# Studying protein-protein interaction through side-chain modeling method OPUS-Mut

**DOI:** 10.1101/2022.05.15.492033

**Authors:** Gang Xu, Yilin Wang, Qinghua Wang, Jianpeng Ma

## Abstract

Protein side chains are vitally important to many biological processes such as protein-protein interaction. In this study, we evaluate the performance of our previous released side-chain modeling method OPUS-Mut, together with some other methods, on three oligomer datasets, CASP14 (11), CAMEO-Homo (65), and CAMEO-Hetero (21). The results show that OPUS-Mut outperforms other methods measured by all residues or by the interfacial residues. We also demonstrate our method on evaluating protein-protein docking pose on a dataset Oligomer-Dock (75) created using the top 10 predictions from ZDOCK 3.0.2. Our scoring function correctly identifies the native pose as the top-1 in 45 out of 75 targets. Different from traditional scoring functions, our method is based on the overall side-chain packing favorableness in accordance with the local packing environment. It emphasizes the significance of side chains and provides a new and effective scoring term for studying protein-protein interaction.

## Introduction

Protein-protein interaction is essential for many biological systems, and it is also important in designing peptidic drugs [1]. Many protein-protein interactions are mediated by amino-acid side chains, especially those of the interfacial residues [2]. Therefore, accurate side-chain modeling for the interfacial residues is crucial.

In recent years, many successful backbone-dependent side-chain modeling methods have been proposed [3-15]. The sampling-based methods select the rotamers from the rotamer library according to their search schemes, and keep the best rotamer for each residue with the minimal score depending on their scoring functions. This kind of methods run fast, and they are suitable for the repeatedly-applied side-chain modeling in the folding process, examples include SCWRL4 [12] and FASPR [14]. However, the performance is limited by the discrete rotamers in the rotamer library and the accuracy of the scoring function [4]. With the help of deep learning techniques, some new methods have been developed, which successfully capture the local environment of each residue, and improve the accuracy of side-chain modeling by a large degree. Examples include DLPacker [5] and OPUS-Mut [15].

Scoring protein-protein docking poses is another important task in studying protein-protein interaction. Various criteria [16] have been proposed, such as force-field based criteria [17, 18], knowledge-based criteria [19, 20], and machine learning-based criteria [21]. Since the side chains of the residues, especially those located at the interface, are crucial for protein-protein interaction, a scoring term that is mainly based on side chains may be an effective term for better scoring the protein-protein docking poses.

In this paper, we use the structures of oligomers to study protein-protein interaction. We evaluate the side-chain modeling performance of our previous released method OPUS-Mut [15], along with some other methods, on three oligomer datasets collected in this study, CASP14 (11), CAMEO-Homo (65), and CAMEO-Hetero (21). The results show that OPUS-Mut outperforms other methods measured by all residues, or by the interfacial residues. To evaluate the performance of OPUS-Mut in scoring protein-protein docking poses, we create a protein-protein docking pose dataset Oligomer-Dock (75) using the top 10 predictions from ZDOCK 3.0.2 [22]. The results show that our scoring function OPUS-Mut (*S*_*all*_) correctly identifies the native pose as the top-1 in 45 out of 75 targets, and ranks native pose among top-3 poses in 67 out of 75 targets. This indicates the effectiveness of using the overall side-chain packing favorableness in accordance with the local packing environment to evaluate different docking poses. In addition, we verify the performance of OPUS-Mut in studying protein mutation on an oligomeric target, SARS-CoV-2 NSP7–NSP8 complex. The results suggest that the usage of OPUS-Mut in studying protein mutation may be generalized to oligomeric complexes.

## Methods

### Datasets

Three oligomer datasets are collected in this study: CASP14 (11) contains 11 oligomers downloaded from the CASP14 website (https://predictioncenter.org/download_area/CASP14/targets), CAMEO-Homo (65) and CAMEO-Hetero (21) contain 65 homo-oligomers and 21 hetero-oligomers, labeled by the CAMEO website [23] released between November 2021 and February 2022, respectively. Note the oligomers with over 3000 residues in length have been excluded from the datasets because of the limitation of GPU memory.

A protein-protein docking pose dataset Oligomer-Dock (75) is created in this study. Among all 97 oligomers from three oligomer datasets, we exclude the oligomers with more than 4 peptide chains, and retain 75 oligomers. For each oligomer, we use the top 10 poses generated by ZDOCK 3.0.2 [22] as its decoys. In the calculation of ZDOCK 3.0.2, for each oligomer, the last peptide chain in the PDB file is defined as “ligand”, the remaining peptide chains are defined as “receptor”.

### OPUS-Mut

OPUS-Mut is a backbone-dependent side-chain modeling method that was released by us recently [15]. It is mainly based on OPUS-Rota4 [4], but with some improvements. The method was shown to outperform some other methods, measured by all residues or by core residues only, on the targets with single peptide chain. In our previous study[15], we use OPUS-Mut to study protein mutation. Briefly speaking, as shown in lower green panel in Figure 1, by comparing the differences between its predicted unmutated (wild-type) side chains and its predicted mutated side chains, we can infer the extent of structural perturbation and the affected residues from those side chains significantly shifted upon the mutation. Also, from the extent of side-chain structural perturbation, we can infer the minimally disturbing mutation, from which we may construct a protein with relatively low sequence homology but with similar structure with respect to the wild type. From the affected residues, we may also use them to infer the possible functional changes if the functions are related to certain residues.

**Figure 1.**
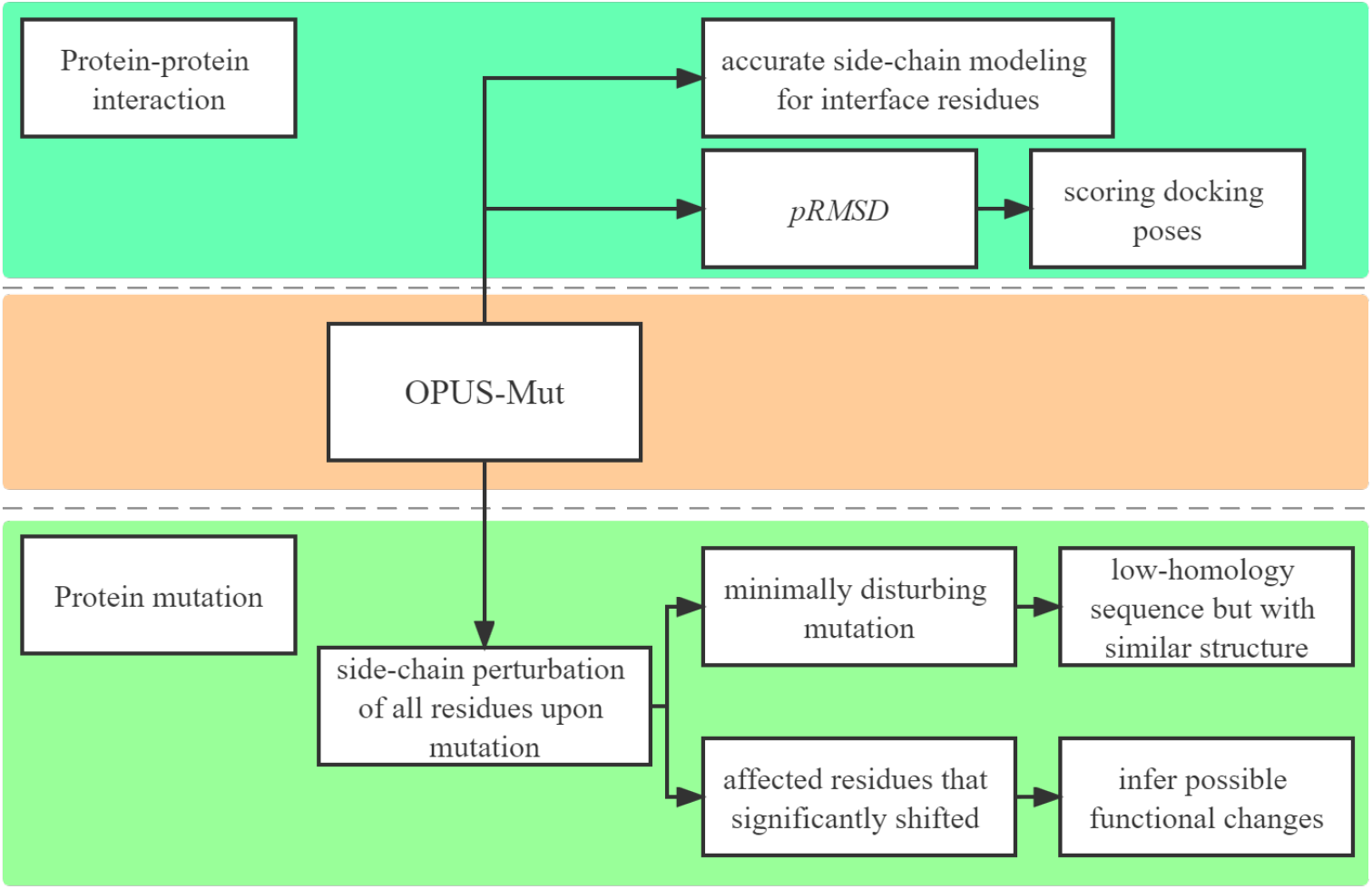
Two applications of backbone-dependent side-chain modeling method OPUS-Mut. The applications of OPUS-Mut in studying protein mutation on the target with single peptide chain (lower green panel) have been evaluated in our previous work. In this paper, we focus on its application in studying protein-protein interaction (upper green panel), and its application in studying protein mutation on oligomeric complexes.

For each residue, besides outputs the predicted side chain conformation, OPUS-Mut also outputs the predicted Root Mean Square Deviation (pRMSD) for its side-chain prediction (upper green panel in Figure 1). To this end, a classification node is used to learn the RMSD between the predicted side chain and its native counterpart for each residue. The pRMSD ranges from 0 to 1, and is segmented into 20 bins. Cross-entropy loss is used for training. In addition, OPUS-Mut adopts a 3DCNN module [5] to capture the local environment for each residue, therefore it can respond to the change of local environment with high sensitivity.

For the residue with lower pRMSD value, OPUS-Mut predicts its side chain with a higher confidence in accordance with its local environment. In this study, we use the summation of pRMSD as an indicator to gauge the overall side-chain packing favorableness in a protein structure, i.e., likeliness of its local packing environment to the native packing environment. In studying docking pose, we name the summation of pRMSD over all residues as *S*_*all*_, the summation over interfacial residues as *S*_*interface*_, and the summation over other residues as *S*_*other*_.

In our pervious study, for the targets with single peptide chain, we have demonstrated that OPUS-Mut outperforms some other backbone-dependent side-chain modeling methods, measured by all residues or by core residues only. In this study, we evaluate the performance of OPUS-Mut and some other backbone-dependent side-chain modeling methods on oligomeric targets. For comparison, we also use OPUS-Mut to model each peptide chain separately, i.e., to model the conformation of side chains without considering the effects of other peptide chains. We name this single-chain approach as OPUS-Mut-s.

### Data availability

The code and pre-trained models of OPUS-Mut and the four datasets used in this paper can be found at http://github.com/OPUS-MaLab/opus_mut. They are freely available for academic usage only.

## Results

### Performance on side-chain modeling on oligomeric targets

We compare the side-chain modeling performance of OPUS-Mut with that of SCWRL4 [12], OSCAR-star [13] and DLPacker [5] on three oligomer datasets CASP14 (11), CAMEO-Homo (65), and CAMEO-Hetero (21). In terms of residue-wise percentage of correct prediction with a tolerance criterion 20° for all side-chain dihedral angles (from *χ*_*1*_ to *χ*_*4*_), OPUS-Mut outperforms other methods measured by all residues (Figure 2a), or by the residues located at the interfaces between different peptide chains (Figure 2b). In this study, the residues with at least one nearby residue (C_α_-C_α_ distance < 10 Å) located at other peptide chain(s) are defined as interfacial residues, and the rest residues are defined as other residues.

**Figure 2.**
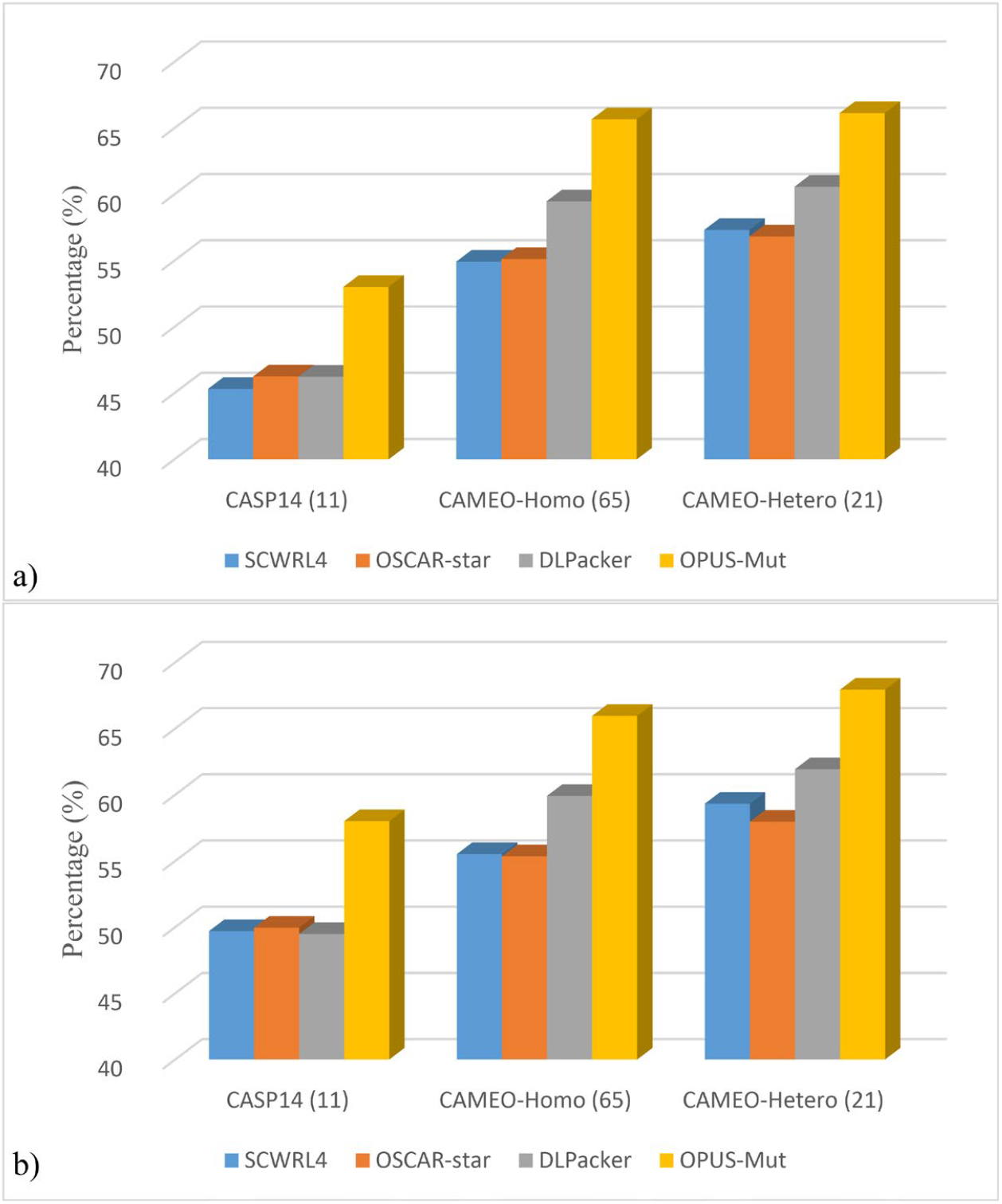
The residue-wise percentage of correct prediction with a tolerance criterion 20° for all side-chain dihedral angles (from *X*_*1*_ to *X*_*4*_) of different methods on three oligomer datasets. a) shows the results measured by all residues. b) shows the results measured by interfacial residues.

As examples, we show two cases of OPUS-Mut side-chain modeling results on interfacial residues and their corresponding experimentally determined crystal structures in Figure 3.

**Figure 3.**
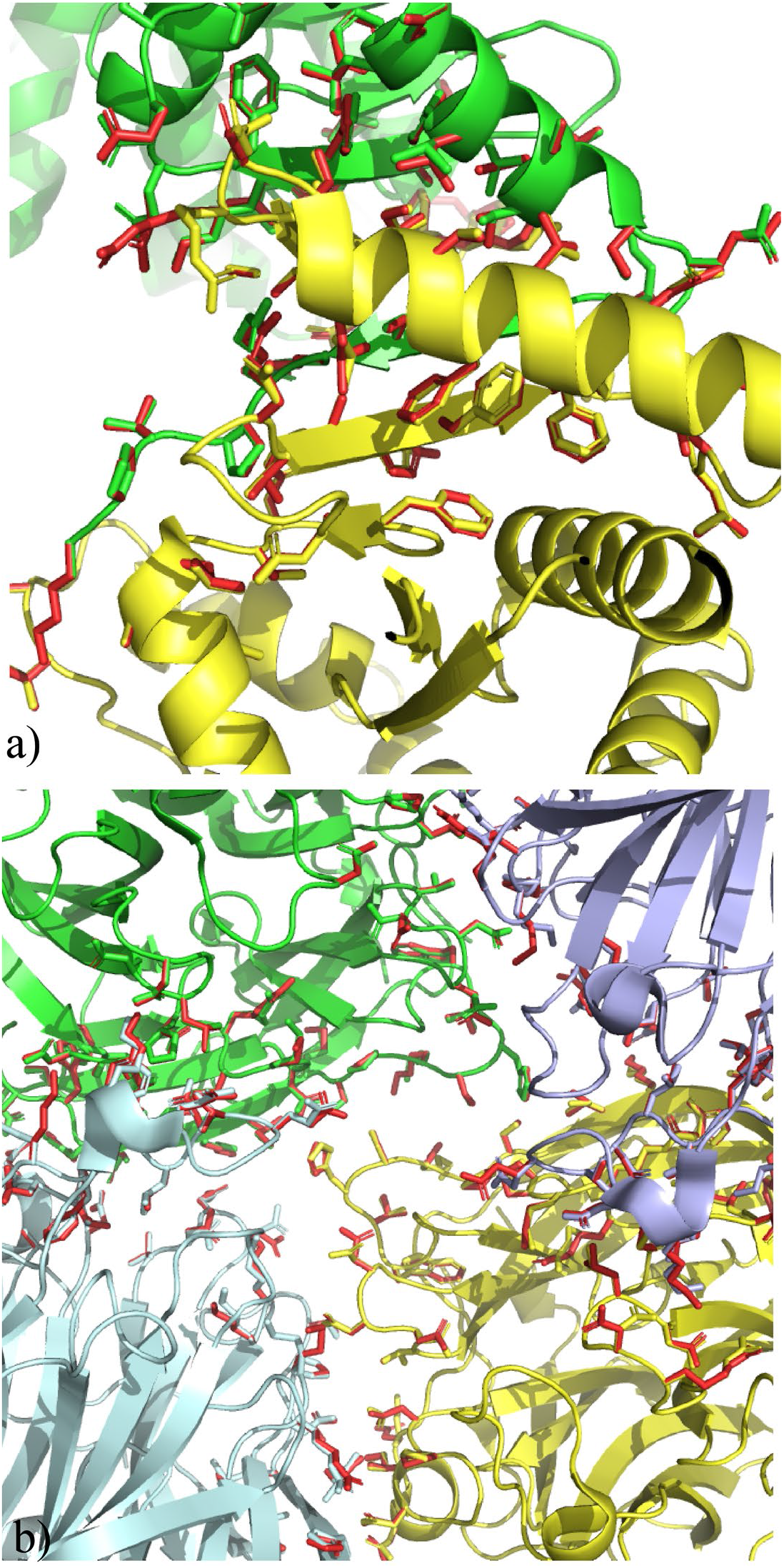
Side-chain modeling results of OPUS-Mut on interfacial residues. The experimentally determined crystal structures are marked in various colors for each peptide chain. The side chains predicted by OPUS-Mut are marked in red. a) and b) show the results of homo-oligomer 7DK9 and 7FIP, respectively.

For a particular peptide chain, to evaluate the influence of other peptide chains on it, we use OPUS-Mut-s, which doesn’t take the effect of other peptide chains into consideration, and models each peptide chain separately. As shown in Table 1, OPUS-Mut outperforms OPUS-Mut-s measured by all residues. While the performance on other residues is almost the same, and the differences are mainly seen in the interfacial residues, for which the performance of OPUS-Mut is significantly better than that of OPUS-Mut-s.

**Table 1.** The residue-wise percentage of correct prediction with a tolerance criterion 20° for all side-chain dihedral angles (from *X*_*1*_ to *X*_*4*_) of OPUS-Mut and OPUS-Mut-s on three oligomer datasets.

|  | All residues | Interfacial residues | Other residues |
| --- | --- | --- | --- |
| CASP14 (11) |  |  |  |
| OPUS-Mut | 53.02% | 58.00% | 49.79% |
| OPUS-Mut-s | 49.36% | 49.18% | 49.48% |
| CAMEO-Homo (65) |  |  |  |
| OPUS-Mut | 65.67% | 65.95% | 65.57% |
| OPUS-Mut-s | 64.17% | 60.17% | 65.60% |
| CAMEO-Hetero (21) |  |  |  |
| OPUS-Mut | 66.14% | 67.93% | 65.26% |
| OPUS-Mut-s | 64.12% | 61.96% | 65.19% |

### Performance on scoring protein-protein docking poses

For studying protein-protein docking poses, we first examine the effect of the partner peptide chain(s) in oligomers. We compare the summation of pRMSD obtained by OPUS-Mut with that obtained by OPUS-Mut-s on three oligomer datasets, the latter doesn’t take the effects of other peptide chains into consideration. As shown in Table 2, the average values of *S*_*all*_ of the targets in each oligomer dataset is lower than that of OPUS-Mut-s, which means the local packing environment used for side-chain prediction in OPUS-Mut is closer to native than that in OPUS-Mut-s. The average values of *S*_*other*_ between OPUS-Mut and OPUS-Mut-s are almost the same, while the average values of *S*_*interface*_ show significant differences, which indicates that protein-protein interaction may bring a more favorable local packing environment to interfacial residues. In addition, among all 97 oligomers in three datasets, OPUS-Mut is lower than OPUS-Mut-s on 88 out of 97 targets in terms of *S*_*all*_, and 94 out of 97 targets in terms of *S*_*interface*_, respectively.

**Table 2.**
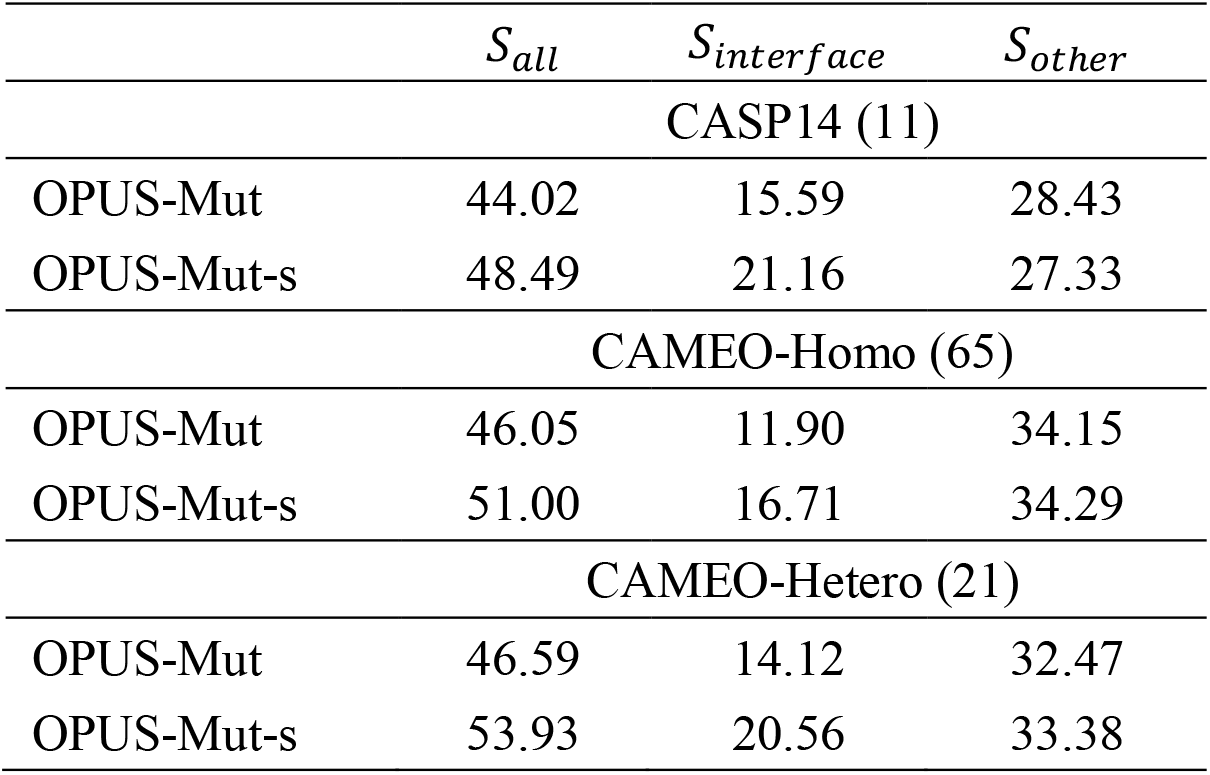
The average values for the summation of pRMSD on three oligomer datasets.

| | $S_{all}$ | $S_{interface}$ | $S_{other}$ |
| --- | --- | --- | --- |
| CASP14 (11) |  |  |  |
| OPUS-Mut | 44.02 | 15.59 | 28.43 |
| OPUS-Mut-s | 48.49 | 21.16 | 27.33 |
| CAMEO-Homo (65) |  |  |  |
| OPUS-Mut | 46.05 | 11.90 | 34.15 |
| OPUS-Mut-s | 51.00 | 16.71 | 34.29 |
| CAMEO-Hetero (21) |  |  |  |
| OPUS-Mut | 46.59 | 14.12 | 32.47 |
| OPUS-Mut-s | 53.93 | 20.56 | 33.38 |

For further investigation, we examine the performance of OPUS-Mut on distinguishing the native docking pose from 10 predicted docking pose decoys for the targets in Oligomer-Dock (75). The results of ZRANK [18] are also listed for comparison. Before the calculation of ZRANK, we use *addh* in Chimera 1.14 [24] to add the hydrogens for each PDB file, then the “TER” line is added between ligand and receptor atom coordinates. As shown in Table 3, using the summation of pRMSD over all residues (OPUS-Mut (*S*_*all*_)) as a scoring function, our method correctly identifies the native pose as the top-1 in 45 out of 75 targets, and ranks native pose among top-3 poses in 67 out of 75 targets. The corresponding results from ZRANK are 16 and 56, respectively. These data indicate that our scoring function, which is based on the overall side-chain packing favorableness in accordance with the local packing environment, is an effective term for scoring protein-protein docking poses.

**Table 3.** Number of targets whose native docking poses can be successfully distinguished from their decoys in Oligomer-Dock (75) by different methods.

| | OPUS-Mut ($S_{all}$) | OPUS-Mut ($M_{interface}$) | ZRANK |
| --- | --- | --- | --- |
| TOP1 | 45 | 47 | 16 |
| TOP3 | 67 | 61 | 56 |

Since the interfacial residues vary in different docking poses, we therefore use 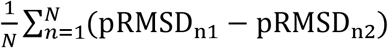, as a scoring function (OPUS-Mut (*M*_*interface*_)), to measure the extent of the improvement of the interfacial side-chain packing favorableness upon docking. For each pose, *N* is the number of interfacial residues, pRMSD_n1_ denotes the pRMSD of the residue *n* predicted by OPUS-Mut-s, pRMSD_*n*2_ denotes the pRMSD of the residue *n* predicted by OPUS-Mut. We assume that a larger *M*_*interface*_ refers to a better docking pose. By using OPUS-Mut (*M*_*interface*_) as a scoring function, as shown in Table 3, our method correctly identifies the native pose as the top-1 in 47 out of 75 targets, and ranks native pose among top-3 poses in 61 out of 75 targets.

As examples, we show docking pose evaluation results on target T1080o in Figure 4. The score of OPUS-Mut (*S*_*all*_) for native pose (Figure 4a) is 7.78, which is the top-1 (lowest) ranking. The scores for other decoy poses are higher, e.g., 8.82 for the pose in Figure 4b, 14.54 for the pose in Figure 4c, and 18.04 for the pose in Figure 4d. Moreover, in Figure 4b, the docking pose is close to the native state (DockQ [25] score 0.948), the results show that the score of ZRANK for this pose is -841.8, lower than the native state score of -793.2, indicating that ZRANK does not identify the correct native pose in this case, while “Ours” does.

**Figure 4.**
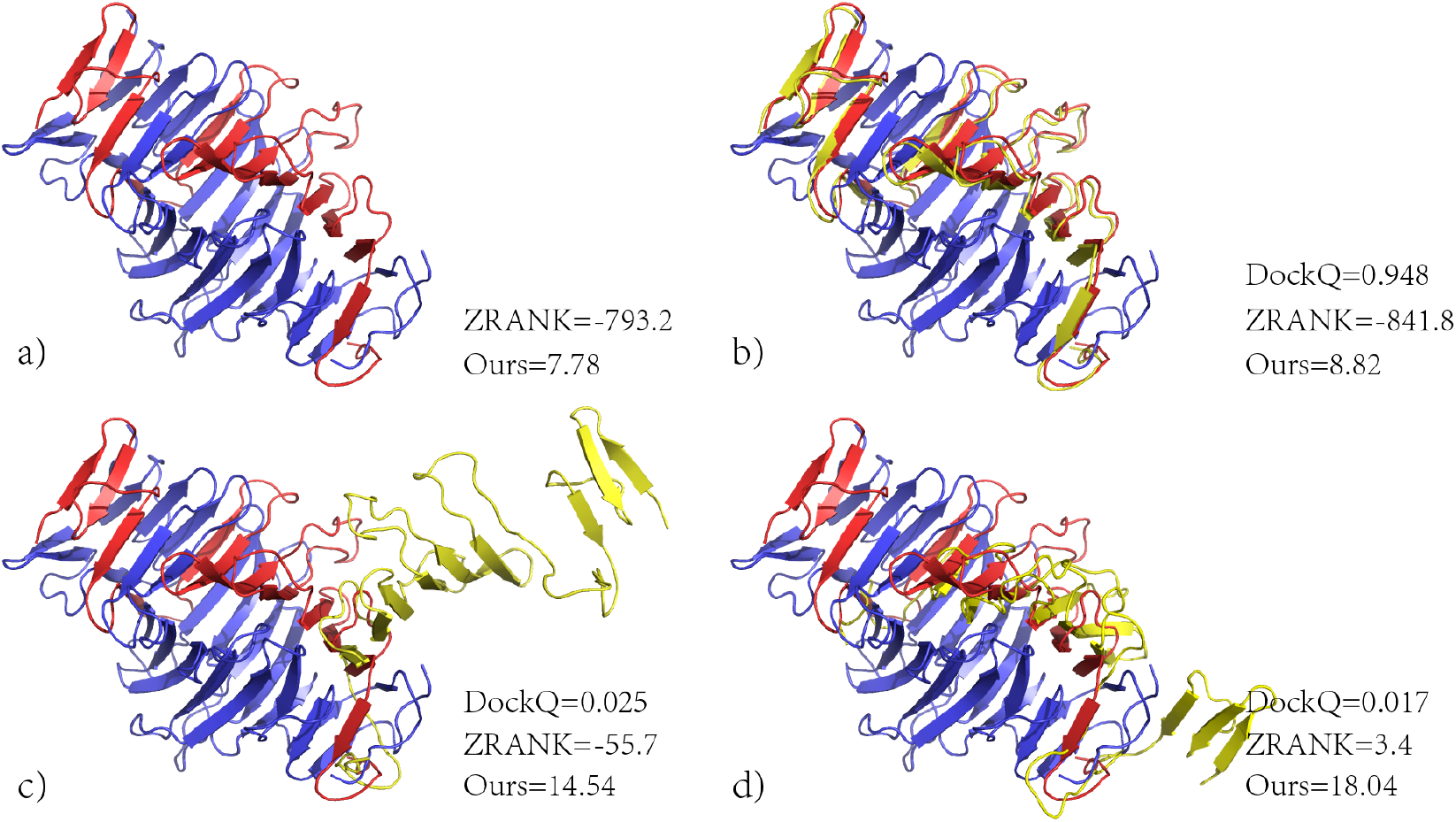
Docking poses evaluation results on T1080o. The structures of native receptor are marked in blue, the structures of native ligand are marked in red. The structures of ligand predicted by ZDOCK 3.0.2 are marked in yellow. We use “Ours” to denote the score from OPUS-Mut (*S*_*all*_). The pose with lower score is closer to the native pose.

### Performance on studying protein mutation on oligomeric targets

Two conserved oligomer interfaces of NSP7 and NSP8 have been studied by Biswal *et al* [26]. NSP7 and NSP8 belong to a complex of non-structural proteins (NSP) that regulates the SARS-CoV-2 RNA-dependent RNA polymerase (RdRP) activity of NSP12. According to Biswal *et al*, NSP7-NSP8 complex is mediated by two distinct oligomer interfaces: interface I responsible for hetero-dimeric NSP7-NSP8 assembly, and interface II mediating hetero-tetrameric interaction between the two NSP7-NSP8 dimers [26]. The interface I contains the following residues: K2, D5, V6, T9, L13, S15, V16, Q19, V66, I68, L71, E74, M75, Q31, F49, K51, V53, S57, L60, and S61 in NSP7, and R80, T84, M87, Q88, T89, L91, F92, R96, L98, N100, L103, I106, P116, I119, I120, and L122 in NSP 8. The interface II contains the following residues: S4, K7, C8, V11, V12, H36, N37, and L40 in NSP7, and V83, M87, M90, T93, and M94 in NSP8. In addition, some residues in interface II of the NSP7–NSP8 tetrameric complex (i.e., S4, C8, V11, V12, N37, and L40 in NSP7, and T84, M87, M90, and M94 in NSP8) are also involved in contacts with NSP12.

According to Biswal *et al* [26], the interface I mutations NSP7^F49A^, NSP7^M52A^, NSP7^L56A^, and NSP8^F92A^ impair the NSP7–NSP8 association. The interface II mutations NSP7^C8G^, NSP7^V11A^, NSP8^M90A^, and NSP8^M94A^ impair the NSP7-NSP8 hetero-tetramer formation. The results also show that, the interface II can not only maintain the hetero-tetrameric assembly of NSP7–NSP8, but also helps to stabilize the hetero-dimeric assembly of NSP7–NSP8. The mutation NSP7^N37V^ does not affect the stability of the NSP7-NSP8 hetero-tetramer appreciably, but it leads to a modest disruption of the NSP7-NSP8-NSP12 complex. According to Subissi *et al* [27], the mutations NSP8^K82A^, and NSP8^S85A^ do not affect the NSP8-NSP12 interaction, but lead to activity loss.

In this study, we download the SARS-CoV-2 NSP7–NSP8 complex (PDB: 7JLT) [26]. Then, we substitute the residues according to the corresponding mutations mentioned above and reconstruct their side chains with OPUS-Mut. Similar to our previous study, we define the affected residue as ones whose mean absolute error of all predicted side-chain dihedral angles (from *χ*_*1*_ to *χ*_*4*_) between the wild-type and mutation is greater than 5 degree. The rest of side chains of other residues are deemed relatively unshifted. All of the affected residues are listed in Table 4 for each mutation.

**Table 4.** Affected residues of different mutations predicted by OPUS-Mut and catalogized by their locations. Experimentally, on the interface I mutations NSP7^F49A^, NSP7^M52A^, NSP7^L56A^, and NSP8^F92A^ impair the NSP7–NSP8 association; on the interface II mutations NSP7^C8G^, NSP7^V11A^, NSP8^M90A^, and NSP8^M94A^ impair the NSP7-NSP8 hetero-tetramer formation. The interface II maintains the hetero-tetrameric assembly of NSP7–NSP8, and also stabilizes the hetero-dimeric assembly of NSP7–NSP8. The mutation NSP7^N37V^ does not destabilize the stability of the NSP7-NSP8 hetero-tetramer significantly, but causes a modest disruption of the NSP7-NSP8-NSP12 complex. The mutations NSP8^K82A^, and NSP8^S85A^ do not affect the NSP8-NSP12 interaction, but result in activity loss.

|  | Interface I | Interface II | NSP8-NSP12 | Others |
| --- | --- | --- | --- | --- |
| NSP7<sup>F49A</sup> | K2, V6, V66,<br>M75 |  |  | M3, T45, M52,<br>S25 |
| NSP7<sup>M52A</sup> | V66, K51 |  |  |  |
| NSP7<sup>L56A</sup> | T9, V66 |  |  | M52 |
| NSP8<sup>F92A</sup> | L71, V66, Q88 |  |  |  |
| NSP7<sup>C8G</sup> | V66 | M90, T93, M94 | M90, M94 | S25, R79, T81 |
| NSP7<sup>V11A</sup> | T9, S15, Q19,<br>R80, M87, M75 | H36, M87 | M87 | R21, E23, M52,<br>D78, S25, Q34 |
| NSP8<sup>M90A</sup> | V66, M75 | T93 |  | S85, T81 |
| NSP8<sup>M94A</sup> | T9, L71 | T93 |  | M52, S85 |
| NSP7<sup>N37V</sup> | T9, Q19, R80,<br>M75, M87 | H36, L40, M87 | L40, M87 | R21, E23, Q34,<br>S25 |
| NSP8<sup>K82A</sup> |  |  |  | E77 |
| NSP8<sup>S85A</sup> | V66, M75 |  |  |  |

As shown in Table 4, according to the results from OPUS-Mut, the mutation NSP7^F49A^, NSP7^M52A^, NSP7^L56A^, and NSP8^F92A^ may cause significant side-chain shift of several residues involved in interface I. Mutations NSP7^C8G^, NSP7^V11A^, NSP8^M90A^, and NSP8^M94A^ may cause significant shift of several residues involved in both interface I and interface II. Mutations NSP8^K82A^ and NSP8^S85A^ may not affect the NSP8-NSP12 interaction although both mutations are close to the residues involved in NSP8-NSP12 interaction (T84, M87, M90, and M94). The predicted results of NSP7^N37V^ are somewhat inconclusive, the experimental results show that NSP7^N37V^ does not affect the stability of the NSP7-NSP8 hetero-tetramer appreciably although it causes a modest disruption of the NSP7-NSP8-NSP12 complex. Our predictions show NSP7^N37V^ may cause significantly shift of several residues involved in interface I, interface II, and NSP8-NSP12 interface, which implies that it could affect the stability of the NSP7-NSP8 hetero-tetramer and the formation NSP7-NSP8-NSP12 complex.

## Concluding Discussion

Protein side chains, especially those located at the interfaces, are crucial for protein-protein interaction. In this study, we study protein-protein interaction through side-chain modeling. We evaluate the performance of several backbone-dependent side-chain modeling methods on three oligomer datasets. The results show that our pervious released method OPUS-Mut [15] outperforms other methods measured by all residues or by the interfacial residues (Figure 2), and its side-chain modeling results for the residues located between different peptide chains are very close to their experimentally determined crystal structures (Figure 3).

When omitting the influence of partner peptide chain(s) in an oligomer and modeling the side chains of each peptide chain separately, the modeling accuracy on oligomer will decrease, especially for that on interfacial residues (OPUS-Mut-s, Table 1). This result indicates that the side chains of interfacial residues may experience conformational changes upon protein-protein association, and it also demonstrates the sensitivity of OPUS-Mut towards local environmental changes.

OPUS-Mut can output the predicted Root Mean Square Deviation (pRMSD) for its predicted side chains on each residue. For a particular residue, lower pRMSD value indicates that OPUS-Mut predicts its side chain with a higher confidence in accordance with its local environment. In this study, we use the summation of pRMSD as an indicator to gauge the overall packing favorableness of side-chain in a protein structure, i.e., likeliness of its local packing environment to the native packing state.

As shown in Table 2, the average values of the summation of pRMSD over all residues (*S*_*all*_) obtained by OPUS-Mut is lower than that obtained by OPUS-Mut-s. The average values of *S*_*other*_ between OPUS-Mut and OPUS-Mut-s are almost the same, while the average values of *S*_*interface*_ show significant differences. Among all 97 oligomers in three datasets, OPUS-Mut is lower than OPUS-Mut-s on 88 out of 97 targets in terms of *S*_*all*_, and 94 out of 97 targets in terms of *S*_*interface*_, respectively. The results indicate that protein-protein interaction may bring a more favorable local packing environment to the interfacial residues.

We compare the performance of identifying native docking pose based on pRMSD from OPUS-Mut with that of ZRANK on Oligomer-Dock (75). Using the summation of pRMSD over all residues (OPUS-Mut (*S*_*all*_)) as a scoring function achieves better results than that using the result from ZRANK, either on correctly identifying native pose, or on ranking native pose in the top three poses (Table 3). Note that, using the summation of pRMSD on interfacial residues as a scoring function (OPUS-Mut (*M*_*interface*_)) may have a bias since the interfacial residues vary in different docking poses. Therefore, we recommend using OPUS-Mut (*S*_*all*_) as a scoring function for scoring different poses. Along with the examples shown in Figure 4, we show that, our scoring function OPUS-Mut (*S*_*all*_), which is based on the overall side-chain packing favorableness in accordance with the local packing environment, may be an effective term for better scoring protein-protein docking poses.

We also verify the performance of OPUS-Mut in studying protein mutation on oligomeric target, SARS-CoV-2 NSP7–NSP8 complex. As shown in Table 4, most of our results are consistent with the experimental results, which indicates that the usage of OPUS-Mut in studying protein mutation may be generalized to oligomeric target.

## Acknowledgements

The work was supported by Shanghai Municipal Science and Technology Major Project (No.2018SHZDZX01), and ZJLab. The work was also supported by National Key Research and Development Program of China (No. 2021YFF1200400).

